# Hearing Ability of Prairie Voles (*Microtus ochrogaster*)

**DOI:** 10.1101/2021.10.07.463519

**Authors:** Emily M. New, Ben-Zheng Li, Tim C. Lei, Elizabeth A. McCullagh

## Abstract

Hearing ability of mammals can be impacted by many factors including social cues, environment, and physical properties of animal morphology. Despite being used commonly to study social behaviors, hearing of the monogamous prairie vole (*Microtus ochrogaster*) has never been fully characterized. In this study, we measure morphological head and pinna features and use auditory brainstem responses to measure auditory capabilities of prairie voles characterizing monaural and binaural hearing and hearing range. Additionally, we measured unbonded male and female voles to characterize differences due to sex. We found that prairie voles show a hearing range with greatest sensitivity between 8 – 32 kHz, robust binaural hearing, and characteristic monaural ABRs. We show no differences between the sexes for binaural hearing or hearing range, however female voles have increased amplitude of peripheral ABR waves I and II and increased latency of wave IV. Our results confirm that prairie voles have both low and high frequency hearing, binaural hearing, and despite biparental care and monogamy, differences in processing of sound information between the sexes. These data further highlight the necessity to understand sex-specific differences in neural processing that may underly variability in behavioral responses between sexes.

**Highlights:**

- Monogamous prairie voles hear across both low and high frequencies.
- Female prairie voles show differences in monaural hearing compared to males.
- There is no difference in binaural hearing or pinna/head size morphology between the sexes.

## 1. INTRODUCTION

The ability to localize sound sources is a critical task for most animals and aids in the ability to forage for food, communicate with conspecifics, mate, and avoid predators among many behaviors. In mammals, spatial locations of sound sources can be approximated using timing and level difference (ITD and ILD respectively) information between the two ears due to head size and pinna morphology (Grothe et al., 2010). Processing of these cues occurs in the auditory brainstem in discrete areas that integrate ITD and ILD information from each ear. Which cues are available to different species, and their hearing ranges, is important to better understand the variability and robustness of the auditory system across taxa.

Prairie voles (*Microtus ochrogaster*) are a species of rodent that have important traits that may impact their sound localization ability and hearing. They are a monogamous rodent with biparental care where both sexes are highly responsive to social and parental cues such as ultrasonic pup vocalizations (Terleph, 2011; Blake, 2012). They occupy both subterranean and terrestrial environments. Many rodents with exclusive subterranean habitats have limited or reduced acoustic spatial ability (Gerhardt et al., 2017) and indeed middle ear structures would indicate that prairie voles should have intermediate hearing ability (based on data from a closely related species *M. arvalis*)(Lange et al., 2004). However, most studies on prairie vole hearing have focused on their vocalizations and social behavior, not on their actual reception of sound and spatial hearing (Stewart et al., 2015).

In this study we report the hearing ability of the monogamous prairie vole using auditory brainstem responses (ABRs) to measure binaural and monaural hearing in this species and between the sexes. We also measure morphology of head size and pinna, which are additional factors in hearing sensitivity due to their limiting in level or timing difference information (Heffner and Heffner, 2008). We hypothesize that due to increased social pressures on male animals, there would be no differences in hearing between the sexes and that prairie voles would have intermediate hearing ability consistent with middle ear anatomical predictions. 73

## 2. MATERIALS AND METHODS

All experiments complied with all applicable laws, NIH guidelines, and were approved by the Oklahoma State University IACUC.

### 2.1 Subjects

All experiments were conducted on lab-reared prairie voles (*M. ochrogaster*) obtained in 2020 from Dr. Tom Curtis’s colony at the Oklahoma State Health Sciences Campus. Both male and female animals were used (N for each experiment listed in the figure legend). Animals were between 180.6 ± 25.5 days old for females and 177.3 ± 27.0 days old for males (not significantly different p = 0.9309). Animals were housed with 1-2 individuals of similar age and sex post weaning (P (postnatal) 21 days) and maintained on a 14:10 (light (6AM) : dark (8PM) cycle). All animals were sexually naïve. Female prairie voles are induced ovulators so there should be no impact of estrus stage on sex differences observed (Dluzen et al., 1981). Animals were tested between 9AM and 3PM during their light cycle. 88

### 2.2 ABR acquisition

ABR recordings were conducted using similar methods as described in (Benichoux et al., 2018; McCullagh et al., 2020). Briefly, animals were anesthetized with a mixture of i.p. ketamine-xylazine (initial-60mg/kg ketamine, 10 mg/kg xylazine maintenance-25 mg/kg ketamine, 12 mg/kg xylazine) and placed on a heating pad in a small sound attenuating chamber (Noise Barriers Lake Forest, IL, USA). Once the animals were unresponsive to a toe-pinch stimulus, subdermal needle electrodes were placed under the skin at the apex (between the ears), reference (nape), and ground (back leg). Two more electrodes were placed behind each ear though data from those channels was not analyzed or presented here. Evoked potentials from the electrodes were amplified by a Tucker-Davis Technologies (TDT, Alachua, FL, USA) RA4LI head stage and TDT RA16PA preamplifier. Potentials were further amplified using a TDT Multi I/O processor RZ5 and recorded using custom Python software. Data were processed using a second order 50 – 3000 Hz filter and averaged across 500-1000 repetitions per condition with 12 ms of recording time. Sound stimuli were generated using a TDT RP2.1 Real-Time processor controlled by custom Python code at a sampling rate of 97656.25 Hz. Sounds were presented to the animal through TDT MF-1 multi-field speakers (1-24 kHz) or TDT EC-1 electrostatic speakers (32-46 kHz) coupled with custom ear bars fitted with Etymotic ER-7C probe microphones (Etymotic Research Inc, Elk Grove Village, IL, USA) for in-ear calibration (Beutelmann et al., 2015). Sound stimuli included transient (0.1 ms) click stimuli and tone bursts (2 ms ± 1 ms on/off ramp) of varying frequencies and intensities. Sounds were presented every 30 ms with an interstimulus standard deviation of 5 ms.

#### 2.2.1 Monaural ABRs

Evoked potentials in response to single ear click stimulation were recorded for each ear independently and quantified as peak amplitude (value from peak to trough) and latency (time to peak amplitude) across the four peaks of the ABR waveform at 90 dB SPL (Figure 1A). Monaural peak amplitude and latency for the left and right ear were then averaged for each animal to obtain a monaural peak amplitude and latency. In addition, click threshold was determined by decreasing the intensity of the sound in 10 dB SPL steps until there was no longer an ABR response recorded. Threshold was then determined to be the difference between the last intensity that elicited a response and the next lowest stimulation (i.e., if 60 dB SPL elicited a response and 50 dB SPL did not, threshold would be determined as 55 dB SPL).

**Figure 1.**
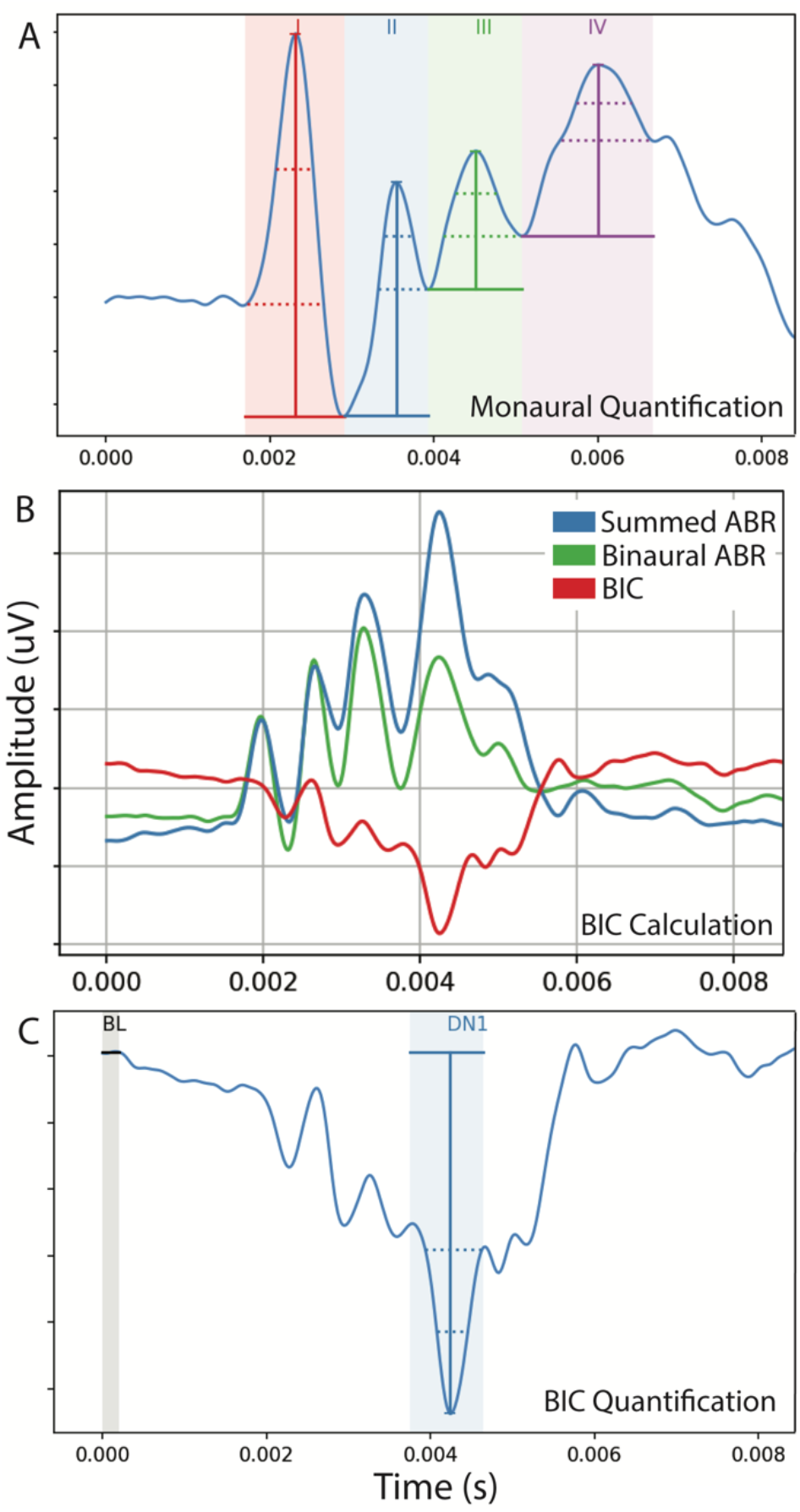
Quantification of monaural/BIC ABRs. Monaural ABRs were quantified as peak to trough (lowest point) for each wave (I, II, III, IV) as shown in A. The BIC was calculated as the binaural ABR subtracted from the sum of the two monaural ABRs at wave IV (B). The BIC amplitude (or DN1) was calculated relative to 0 in the trace (BL or baseline) which was normalized across the entire recording (C).

#### 2.2.2 Audiogram

Threshold, intensity in which there is no longer a neural response, of ABR was determined across frequencies (1, 2, 4, 8, 16, 24, 32, 46 kHz) to generate an audiogram of prairie vole hearing ranges. Briefly, animals were presented with sound intensities in 10 dB steps across frequencies. When a response was no longer observed, threshold was determined to be the difference between the last response and the subsequent lack of response (see above for click threshold).

#### 2.2.3 BIC with ITD

In addition to monaural stimulation, animals were presented with clicks to both ears simultaneously or with an interaural timing difference (ITD, -2 to +2 ms in 0.5 ms steps) at 90 dB SPL. The two monaural responses were summed and subtracted from the binaural ABR response to generate the binaural interaction component (BIC)(Ferber et al., 2016; Laumen et al., 2016; Benichoux et al., 2018, Figure 1B). BIC was then measured across ITDs for both amplitude and latency. The amplitude of the BIC was determined as the peak relative to zero and baseline of the overall trace, which was set to zero to account for variability in the trace (Figure 1C)

### 2.3 ABR data analysis

ABR traces were analyzed using custom Python software. Raw traces (monaural, binaural, BIC) were subtracted by the mean of the initial baseline interval to account for fluctuation of the baseline and zeroing of the baseline across traces. ABR and BIC peaks were automatically detected and labeled by searching local extrema on the detrended trace. The detected peaks were then inspected and manually adjusted if inaccurate. Additionally, if a peak was determined to be not present, it could be deselected and not quantified. Values for amplitude and latency were then saved as separate files and collated across animals.

### 2.4 Morphological characteristics

Morphological features were determined for each animal using 6 Inch Stainless Steel Electronic Vernier Calipers (DIGI-Science Accumatic digital caliper Gyros Precision Tools Monsey, NY, USA). Specifically, pinna size (length and width) and effective diameter (square root of pinna length x pinna width) were measured as well as interaural distance and nose to pinna length. Lastly, animals were weighed using a digital scale.

### 2.5 Statistical analyses

Figures were generated in R (R Core Team, 2013) using ggplot2 (Wickham, 2016) and represent mean and standard error in combined animal line plots. Data were analyzed using a linear mixed effects model to account for repeat observations within one animal (lme4 (Bates et al., 2015)) with sex and condition (ITD, Frequency, Peak) as fixed effects, and animal as a random effect. Linear regression models with sex as an interaction variable were performed for relationships between morphological features and/or ABR waveforms. Two-tailed t-tests were performed to determine if there were differences in age between the sexes and click thresholds. Where values are indicated as statistically significant between the two genotypes, * indicated a p-value of < 0.05, ** = p < 0.01, and *** = p < 0.0001. Figures were prepared for publication using Adobe Illustrator (Adobe, San Jose, CA).

## 3. RESULTS

We measured the hearing ability of prairie voles using auditory brainstem responses and morphological measures. Specifically, ABR responses (amplitude and latency) to monaural click stimuli, ABR threshold across frequencies, BIC of the ABR variability with ITD (amplitude and latency), and morphological features (pinna size, and head characteristics) were measured between the sexes.

### 3.1 Monaural ABRs

We first measured male and female prairie vole’s responses to monaural transient click stimuli (Figure 2). Prairie voles showed a characteristic ABR like other small rodents with four prominent peaks (I-IV, Figure 1A). These peaks had similar latencies to other species, appearing within the first 6-8 ms after the click stimulation (Benichoux et al., 2018). Studies have shown that in humans there are clear differences in amplitude and latencies of speech-ABRs, and generally shorter latencies across ABR-type (Watson, 1996; López-Escámez et al., 1999; McFadden et al., 2010; Liu et al., 2017). There were clear sex differences in monaural peak and amplitude of ABR responses in prairie voles. Peak amplitude was significantly larger in female prairie voles for waves I and II of the ABR compared to males (Figure 2A – individual responses, Figure 2C mean responses). Interestingly, latency of peaks was significantly longer in females for wave IV of the ABR (Figure 2B – individual responses, 2D mean responses). There was no significant difference in overall hearing threshold between male and female prairie voles (as seen by no difference in click threshold responses, data not shown).

**Figure 2.**
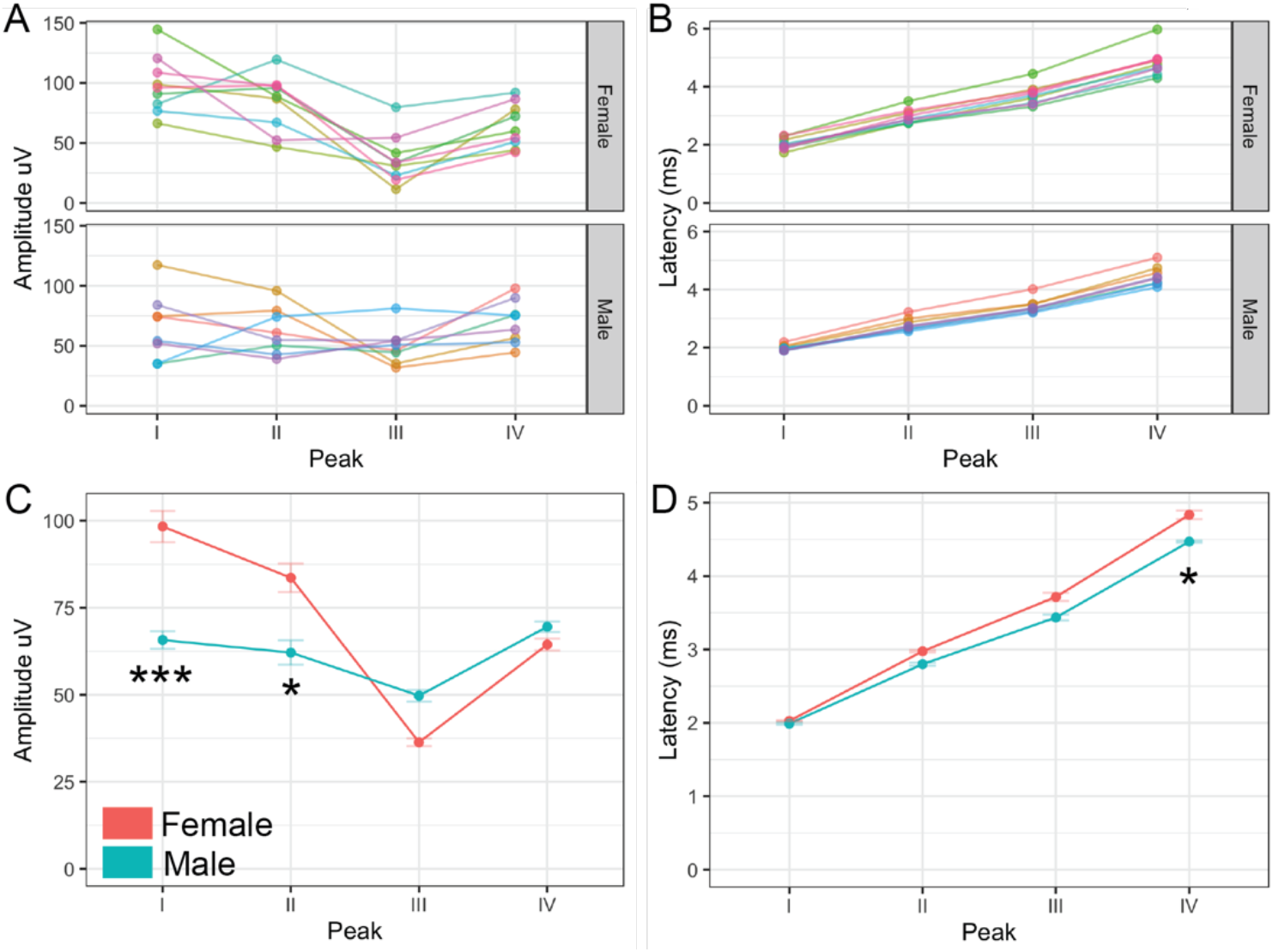
Monaural ABRs. Monaural ABRs were measured for each of the four peaks of the prairie vole ABR. Data for individual animals (average of left and right ABRs) across peak amplitude (A) and latency (B) are shown for male and female animals. The average responses for all male (blue) and female (red) prairie voles for amplitude (C) and latency (D) across peaks I-IV. Data represent 8 males and 9 females * = *p* < *0.05*, *** = *p* < *0.001*

### 3.2 Audiogram

Next, we wanted to determine sex differences in frequency responses. Like the click threshold, there was no difference between the sexes across frequencies (Figure 3A – individual responses, Figure 3B mean responses). Prairie voles show a broad hearing range, with best hearing between 8-32 kHz like other rodents (Lin et al., 2021).

**Figure 3.**
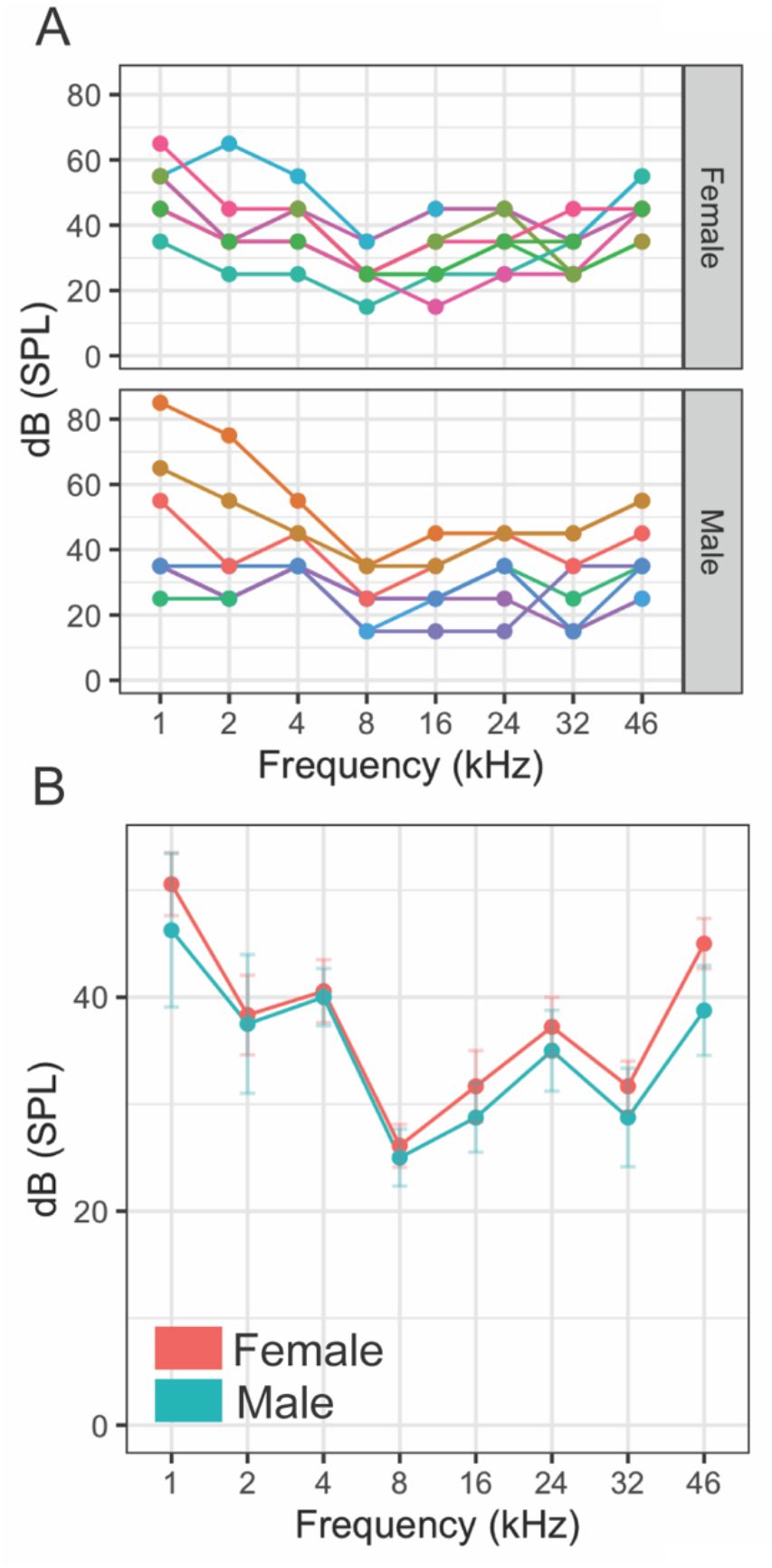
Audiogram. Hearing threshold was measured across frequencies for male and female prairie voles. Data for individual animals across frequency and sex are shown in A (each color/line represents data from one individual). Mean responses for the sexes (red = female, blue = male) show no differences between the sexes (B). N = 8 males and 9 females.

### 3.3 BIC x ITD

Binaural hearing ability of prairie voles was assessed using the binaural interaction component (BIC) of the ABR as it varies with ITD. The BIC has been shown to be a predictive biomarker of binaural hearing ability in many species and is responsive to hearing-impairment (Laumen et al., 2016; Benichoux et al., 2018). Unlike monaural ABRs, there was no difference between the sexes in either amplitude (Figure 4A for individual responses, Figure 4 C for mean responses) or latency (Figure 4B for individual latencies, Figure 4D for mean latencies) of the BIC as it varied with ITD. Like other studies, the BIC amplitude was largest and BIC latency shortest around 0 ITD. BIC latency increased with longer ITDs while amplitude decreased as ITD increased (Ferber et al., 2016; Benichoux et al., 2018).

**Figure 4.**
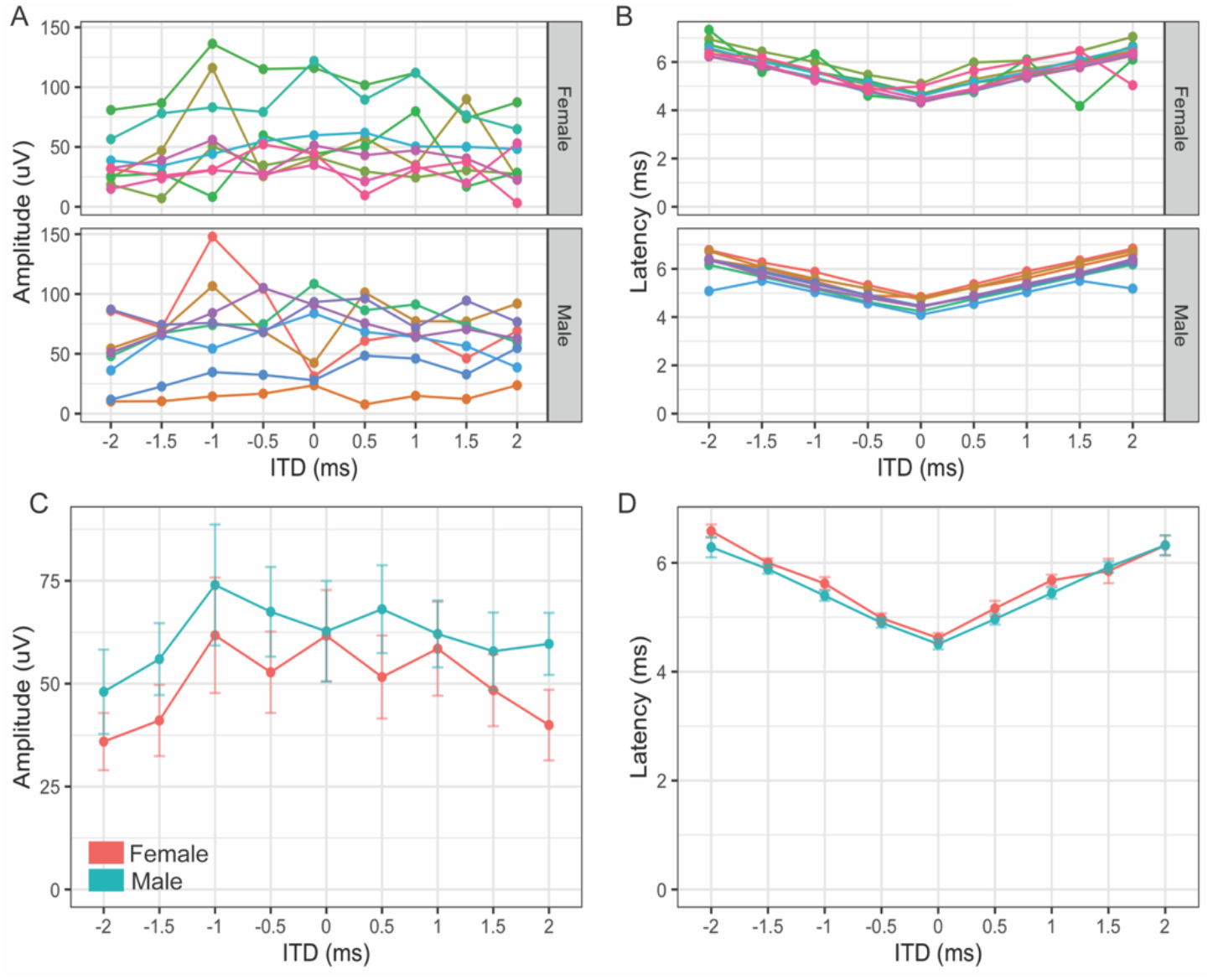
BIC x ITD. The BIC as it varied with ITD was measured for both sexes. Data for individual animals across ITD and sex are shown in A for amplitude and B for latency (each color/line represents data from one individual). Mean responses for the sexes (red = female, blue = male) show no differences between the sexes for either amplitude (C) or latency (D). N = 8 males and 9 females.

### 3.4 Morphology

Pinnae are the first anatomical feature to play a role in hearing ability of an animal with external ears (Butler, 1975). We measured pinna size (length and width), effective diameter (Greene et al., 2014), distance between the pinna (inter pinna length), and nose to pinna distance between the sexes (Figure 5A). We did not observe any statistically significant differences between the sexes for any of the morphological features or comparisons to ABR waveforms (Figure 5). Unsurprisingly, pinna length and width were positively correlated indicating ears grew larger symmetrically (Figure 5B). Similarly inter pinna length and nose to pinna length (Figure 5C), effective diameter of the pinna and inter pinna length (Figure 5D), and weight and effective diameter (Figure 5E) were positively correlated indicating that these attributes generally varied with overall increase of size of individuals. Lastly pinna size in female voles was negatively correlated with Wave I of the ABR while males showed a positive correlation (Figure 5F). While not significant, these data show that pinna size is a possible contributing factor to sex differences seen in ABR Wave I amplitude.

**Figure 5.**
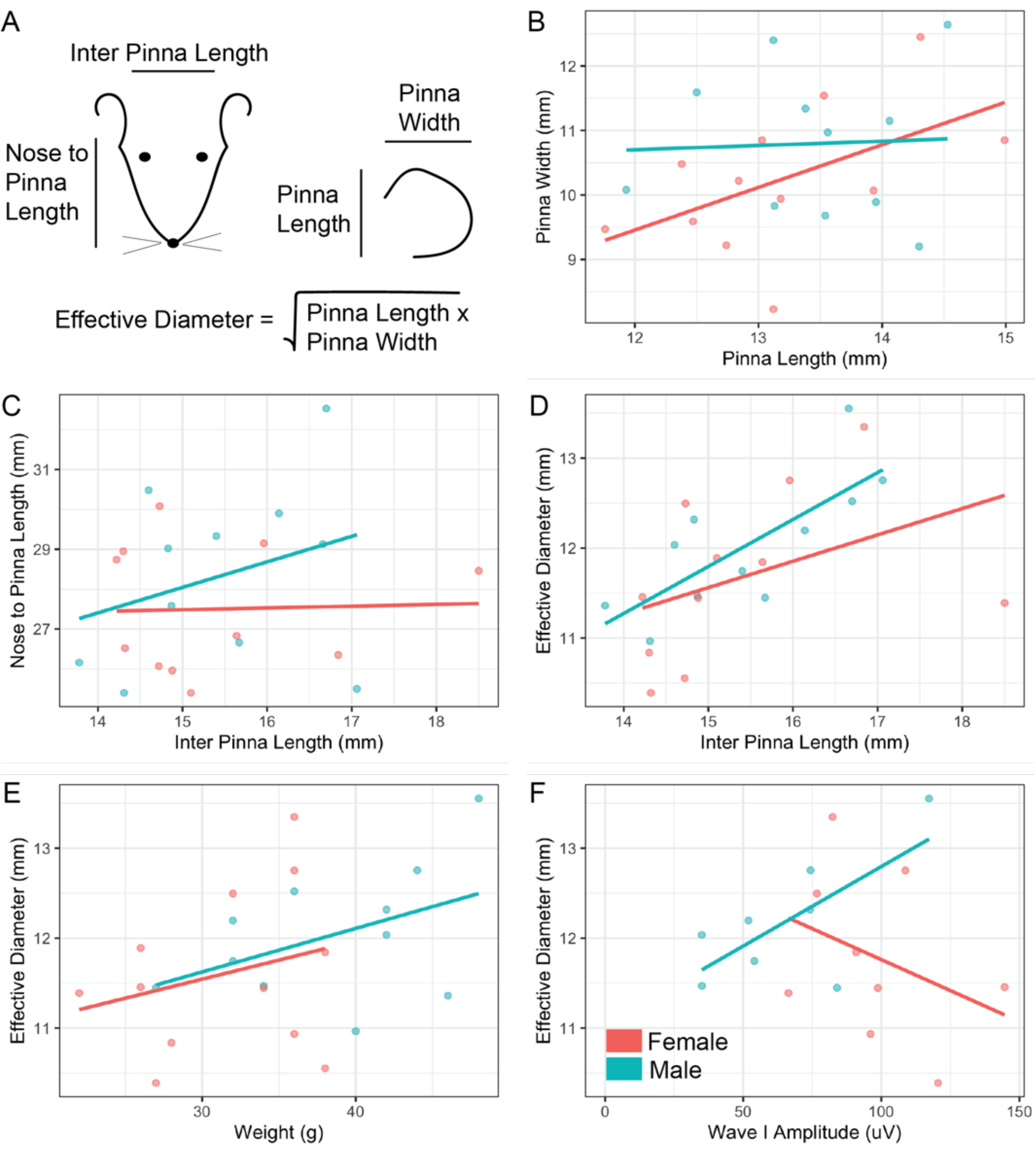
Morphological features. Measurements of anatomical features of the head were taken for both male and female prairie voles (A). Pinna size (B), head features (C), pinna diameter and inter pinna length (D), weight and pinna diameter (E), and pinna diameter and wave I of the ABR (F) were compared (red = females, blue = males). Each dot represented data from one individual. N = 11 males and 12 females.

## 4. DISCUSSION

This is the first study to fully characterize the hearing ability of prairie voles, including both monaural and binaural hearing, frequency range, and pinna morphology between the sexes. While we did not find significant differences across hearing range or binaural hearing ability between the sexes, we did see significant differences in monaural amplitude and latency indicating sex-specific changes in auditory processing in this species. Prairie voles are particularly interesting rodents due to their monogamous breeding structure, where most studies have focused (McGraw and Young, 2010). Hearing ability is an important aspect of social communication that has been understudied in this group. Our hearing range findings are consistent with work on other vole species (Lange et al., 2004) and proposed hearing ranges based on vocalizations (Terleph, 2011; Stewart et al., 2015). However, this is one of the few studies to show differences in amplitude and latency of monaural ABRs between the sexes, particularly in a rodent model that more closely resembles humans in terms of biparental care.

Most of the work regarding sex differences in rodent ABRs shows differences in overall hearing threshold between the sexes (reviewed in Lin et al., 2021). Additionally, sex differences have been observed in several clinical models of hearing disorders in several rodent species or strains (Lin et al., 2021), again mostly as changes in overall ABR or hearing threshold or range. In contrast, our study did not show any differences in either click threshold or hearing range (audiogram) between the sexes. In addition, we observe differences in monaural hearing ability, but no change to binaural hearing ability (as measured by the BIC). Consistent with no changes in binaural hearing, the monaural sex differences we see are increased amplitude of waves I and II in females, which are not thought to be part of the binaural pathway (Alvarado et al., 2012). However, we do show increased latency of wave IV of the ABR in females, which is thought to represent binaural areas. Interestingly, increased latency of responses in females is opposite to sex differences in ABR latencies in humans which are typically shorter in women (López-Escámez et al., 1999; McFadden et al., 2010). Further study exploring sex differences in hearing after mate-pairing in this species would be particularly interesting to study the impact of hormones on hearing.

The choice of rodent model for studying hearing is often limited to genetically tractable models such as rats or mice despite behavioral and hearing range differences to humans. However, in comparison to other neuroscience fields, auditory function has been measured in many different species, in particular rodents (Heffner and Heffner, 2007). Compared to other rodents, prairie voles are somewhat unique in their monogamous mating style, which adds to the breadth of social behavioral comparisons to humans. In addition, since female prairie voles are induced ovulators, estrus stage does not need to be quantified when measuring virgin females unexposed to males post weaning (Dluzen et al., 1981). Despite this, most research on prairie vole communication has focused on vocalizations and not on neural properties of receiving sound information (Stewart et al., 2015; Kolacz et al., 2018). Prairie voles vocalize between 2.5 – 35 kHz and physics of their middle ear would indicate optimal hearing ranges between 20 - 40 kHz which is somewhat consistent with our findings of best hearing between 8 - 32 kHz (Lange et al., 2004; Kolacz et al., 2018). The fact that there are no differences between males and females in ABR-measured hearing range may well be consistent with a monogamous rodent that has biparental care because both males and females respond to pup ultrasonic vocalizations (Terleph, 2011; Blake, 2012).

Our results indicate no sex differences in pinna or head size in prairie voles. Rodents have been postulated to hear high frequencies due to the limitation in their small head size to create interaural level difference cues (Heffner et al., 2001). However, rodents also have some of the broadest hearing ranges of any mammal (Heffner and Heffner, 2008). An important aspect of hearing ability and sound localization is the size and shape of the head and pinna of an animal. Pinnae have an important role in vertical sound localization and removal of front-back confusion (Heffner et al., 1996). Many species have notable secondary peaks of high frequency hearing and prairie voles are no exception, with a secondary peak around 32 kHz. These secondary peaks can be attributed to pinna size, shape, and directionality, with the smaller the size of pinna and head, typically the higher frequency hearing (Heffner and Heffner, 2018). Compared to *Mus musculus* or other similar sized species, prairie voles have small pinna for their overall size, which would indicate that they should hear even higher frequencies. It is possible that prairie voles can hear even higher than the measured 46 kHz or that there is specific directional information that is missing in pure anatomical measurements. In addition, we did not measure enough frequencies in the low hearing range to test the true lower limit of their low-frequency hearing. Presence of a robust medial superior olive (MSO), an important structure for mammalian low frequency hearing, would indicate that they likely have fairly good low frequency hearing as well (unpublished data) making them more similar in hearing range to the Mongolian gerbil (*Meriones unguiculatus*) than the house mouse (*M. musculus*) (Grothe et al., 2010).

## 5. CONCLUSIONS

This is the first study to measure the hearing ability of the monogamous rodent, *Microtus ochrogaster* including morphology, and monaural and binaural stimuli. Importantly, we measured whether there were sex differences in hearing range, morphology, and monaural or binaural responses. We hypothesized, due to the biparental care and monogamy in this species, there would be limited sex differences in hearing ability. We showed that while there were no differences in the sexes for morphological measures, hearing range, and binaural measurements there were significant differences in monaural amplitudes of peripheral auditory brainstem waves (I and II) and increased latency in wave IV of females. These findings are significant because they indicate that in some species, sex differences may be limited to certain aspects of hearing and not others. The mechanisms that underly these differences in amplitude and latency of monaural responses are an intriguing area for future research and may provide insights into hearing differences observed in other species.

## 6. ACKNOWLEDGEMENTS

We would like to acknowledge and thank Dr. Tom Curtis for sharing his prairie vole colony with us and helping us to establish and care for our new colony. We would also like to thank several undergraduate researchers including Jordyn Sisovsky and Adam Ramirez who helped with the early collection of data for this project. We also recognize funding from the Oklahoma State University College of Arts and Sciences Summer Research Award to Dr. McCullagh allowing her to dedicate time and effort to this project over the summer 2021. Lastly, Emily Margaret New is supported through an Oklahoma State University Wentz Research grant for 2021-2022 to conduct undergraduate research in the McCullagh lab.

## 7. AUTHOR CONTRIBUTIONS

EMN: data collection and curation BZL: software, validation, TCL: software, validation EAM: conceptualization, methodology, original draft preparation, visualization, supervision. All authors contributed to reviewing and editing the manuscript.

